# Geometry and evolution of the ecological niche in plant-associated microbes

**DOI:** 10.1101/836411

**Authors:** Thomas M. Chaloner, Sarah J. Gurr, Daniel P. Bebber

## Abstract

The ecological niche of a species can be conceptualized as a volume in multidimensional space, where each dimension describes an abiotic condition or biotic resource. The shape and size of this volume strongly determines interactions among species and influences their global distribution, but the geometry of the niche is poorly understood. Here, we analyse temperature response functions and host plant ranges for hundreds of fungi and oomycetes. We demonstrate that niche specialization is independent on abiotic and biotic axes, that host interactions restrict fundamental niche breadth to form the realized niche, and that both abiotic and biotic niches show limited phylogenetic constraint. Such niche adaptability makes plant pathogens a formidable threat to agriculture and forestry.

## Main text

The niche is a fundamental concept in ecology and evolution, describing the range of conditions under which an organism can survive and reproduce (*1*). Hutchinson’s model of the niche as a volume in multidimensional space (*2*), where each dimension represents an environmental condition or resource requirements affecting a species, has proven a powerful tool for understanding competition, trait evolution, ecological specialization, community assembly rules, and the distributions of species on Earth (*3, 4*). In the era of anthropogenic habitat modification, climate change, and invasive species, modelling the ecological niche is key to predicting and mitigating the impacts of human activities on the biosphere (*5*).

Niche theory differentiates between niche axes defined by abiotic conditions such as temperature or soil pH and biotic resources like host or prey availability. Abiotic conditions are unaffected by the species while resources can be depleted and competed over with other species, resulting in exclusion of the inferior competitor (*3, 6*). Biotic interactions thereby modify our expectations of where a species could exist in nature, reducing and altering the shape of the realized niche in comparison with the fundamental niche (*1*). Details of the geometry of the niche remain unresolved (*4*), such as the shape of the response of metabolic rates to temperature (*7, 8*). While the importance of biotic interactions on the abiotic niche and consequently on species’ geographical distributions is increasingly recognized (*9–11*), there has been little theoretical or empirical research on the relationship between abiotic niche and biotic niche axes. For example, does specialization on abiotic niche axes correlate with specialization on biotic niche axes (*12*)?

Here, we analyse temperature response functions and host ranges of hundreds of plant-associated fungi and oomycetes, to understand the shape and size of the abiotic niche and test whether abiotic and biotic niches are correlated or independent. Most biogeographical and niche modelling studies have been conducted on plants, vertebrates and insects for which distributional data are available at high spatial resolution, and for which important biotic interactions, such as host or prey species, are known (*11, 13–15*). Much less is known about the niche dimensions of microbes, which constitute most biodiversity (*16*) and are key to ecosystem function (*17*). For example, how do temperature responses vary among microbial species (*18*), how does the temperature niche evolve (*19*), and what is the relationship between abiotic and biotic niche axes?

We collated and analysed experimentally-derived temperature responses, specifically the minimum (T_min_), optimum (T_opt_) and maximum (T_max_) temperatures that comprise the ‘cardinal temperatures’, of various biological processes for 661 plant-associated microbes (599 fungi and 62 oomycetes) (Fig. 1). Previous analyses of thermal responses have considered only a handful of fungi and no oomycetes (*20–22*). Cardinal temperatures can be used to derive temperature response functions, or thermal performance curves (*8*), using mathematical forms such as the beta function (*23*). The biological processes for which cardinal temperatures have been measured vary in their degree of host interaction. Experimental measurements for rates of growth in culture (GC) and often spore germination (SG) occur under axenic conditions, while infection (IN) and disease development (DD) occur as interactions with the host plant. Fruiting body formation, or fructification, (FR) and sporulation (SP) may or may not be measured *in planta* depending on experimental conditions. Variation in host interaction among biological processes allows the effect of biotic interactions on the temperature niche to be quantified.

**Fig. 1.**
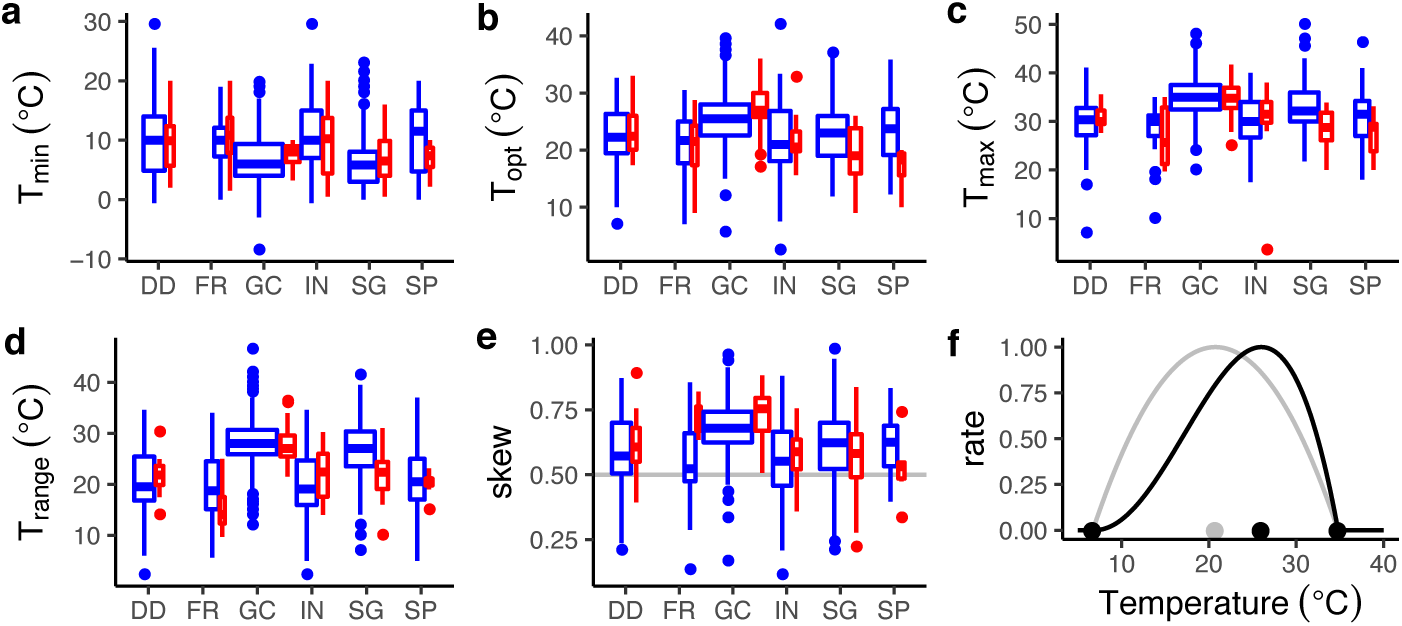
Temperature responses of plant pathogenic fungi (blue) and oomycetes (red) life history processes. A. Minimum temperature, B. Optimum temperature, C. Maximum temperature, D. Temperature range (T_max_ – T_min_). E. Skew, where values >0.5 indicate the T_opt_ is closer to T_max_ than T_min_. The grey line indicates where T_opt_ lies half-way between T_min_ and T_max_. Processes are disease development (DD), fructification (FR), growth in culture (GC), infection (IN), spore germination (SG), and sporulation (SP). Box widths are proportional to the square root of sample size. F. Illustration of skew for a temperature response function. The black points show cardinal temperatures, with midpoint of T_min_ and T_max_ in grey. Unskewed response in grey, skewed response in black.

We found substantial overlap in the distributions of cardinal temperatures between fungi and oomycetes for all processes (Fig. 1, Table S1). GC and SG had somewhat lower T_min_ and higher T_max_ (and hence wider T_range_) than other processes (Table S1). Rates increase with temperature to T_opt_ following thermodynamic expectations, followed by rapid decline in rate as enzymes denature (*8*). We defined asymmetry, or skewness, of the temperature response function as the degree to which T_opt_ is closer to T_max_ (skew > 0.5) or T_min_ (skew < 0.5). For nearly all processes in both fungi and oomycetes, T_opt_ was closer to T_max_ than T_min_, but most strongly for GC (Fig. 1D). This suggests a fundamental difference in the shape of the temperature response for growth in axenic culture (GC) than for processes that involve interaction with the host plant or occur without nutrient media.

Within species, GC and SG tended to have similar cardinal temperatures (Fig. 2, Table S2). GC and SG had lower T_min_ than the other biological processes, higher T_opt_, higher T_max_, a wider T_range_, and greater skew (Fig. 2). T_opt_ values were largely correlated across biological processes (Pearson correlation > 0.5 between most processes, Table S3), but T_range_ values were either poorly, or not significantly, correlated (Table S4) other than between DD and IN (Pearson correlation 0.94, 95% confidence interval 0.89 – 0.97). Species are therefore warm or cold-adapted across biological processes, but there is less evidence that temperature niche breadth is correlated across processes. In summary, T_opt_, T_range_ and skew of the temperature response function were significantly greater for GC and SG than for other processes (Fig. 2E). This phenomenon has been detected in temperature-adapted strains of a fungal species (*24*). The narrower temperature responses for *in planta* processes compared with *in vitro* processes could demonstrate the modification of the Fundamental Niche by biotic interactions to give the Realized Niche (*1*). GC occurs under controlled axenic conditions with optimal nutrient availability, and in the absence of competition or other biotic interactions. Processes relating to disease *in planta* occur in the presence of plant host defences or stress responses and under nutrient restriction compared with processes occurring in culture media, and so can be considered sub-optimal for the pathogen. These suboptimal resource conditions appear to restrict temperature niche breadth, reducing T_opt_ by reducing the relative growth rate at higher temperatures (Fig. 2E). The left-skew of temperature response functions means that reduction in relative rates at high temperatures is much larger than at low temperatures (Fig. 2F).

**Fig. 2.**
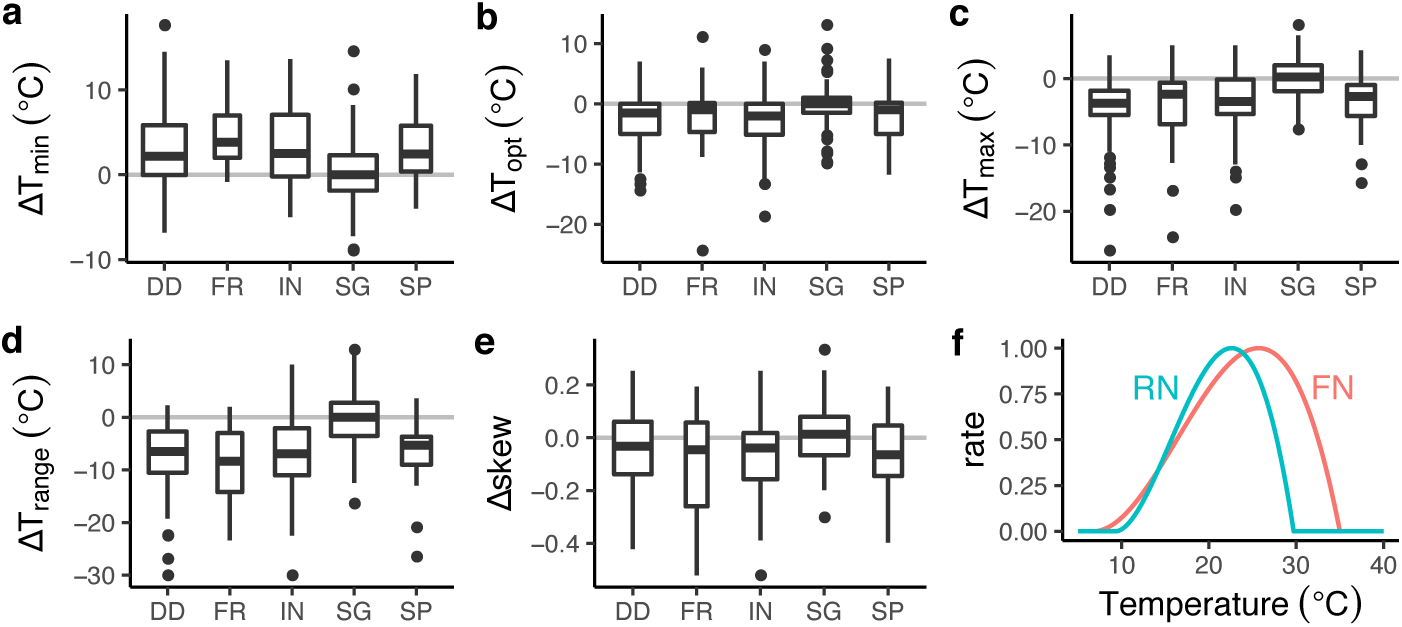
Temperature response differences to growth in culture. A. T_min_, B. T_opt_, C. T_max_, D. T_range_. E. Skew. Processes are disease development (DD), fruitification (FR), infection (IN), spore germination (SG) and sporulation (SP). F. Illustration of temperature response (beta function) for the fundamental niche (red, represented by growth in culture) compared with the realized niche (blue, represented by disease development), where RN has a narrower T_range_ and lower T_opt_ than FN.

Species that occupy relatively large volumes of niche space are commonly described as generalists, while those with narrow tolerances are termed specialists (*12*). There is little empirical understanding or theoretical consideration of the correlation between niche breadth on different niche axes, i.e. is the n-dimensional hypervolume an n-sphere or a hyperellipsoid? We found no evidence for correlation between phylogenetic diversity of known host plants and T_range_, indicating that specialization can occur independently for biotic resources and abiotic conditions (Fig. 3, Table S5). The terms specialist and generalist therefore cannot be applied as holistic descriptions of fungal or oomycete species’ ecology, but it is difficult to speculate on the selection pressures that would lead to differential specialization on abiotic and biotic niches.

**Fig. 3.**
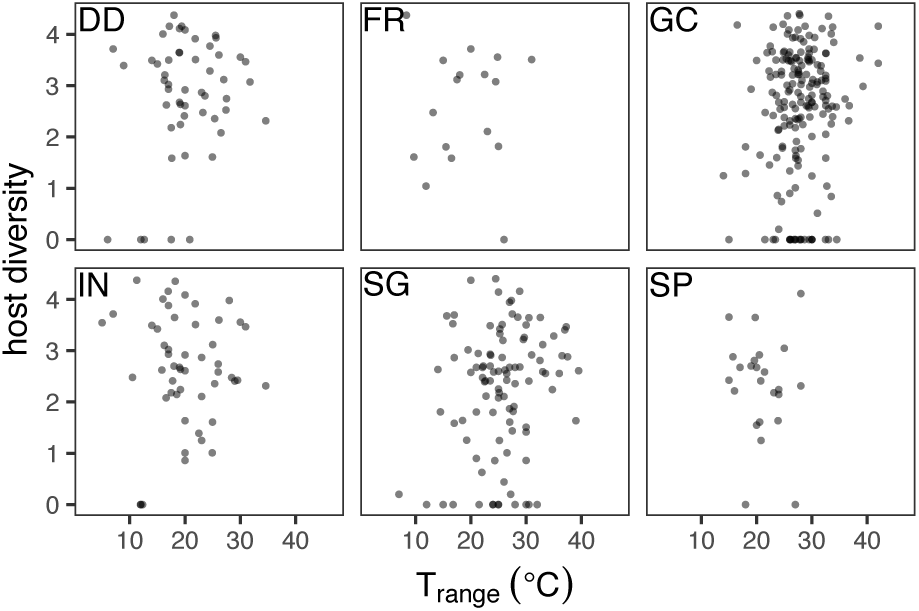
Biotic vs. abiotic niche breadth. Biotic niche breadth is represented by 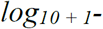 transformed host phylogenetic diversity calculated from the processed host phylogeny, and abiotic niche breadth by T_range_. Panels show the different biological processes, disease development (DD), fructification (FR), growth in culture (GC), infection (IN), spore germination (SG) and sporulation (SP).

We investigated the evolution of abiotic and biotic niche axes via phylogenetic analyses of thermal physiology and host range in the oomycete genus *Phytophthora*. We selected *Phytophthora* because this was the only multi-species genus in the dataset for which well-resolved molecular phylogenies and host range data were available (*26*). We found a small but significant phylogenetic signal in T_opt_ for *Phytophthora* species (Bayesian phylogeny: Blomberg’s *K* = 0.220, p < 0.05; Maximum likelihood phylogeny: Blomberg’s *K* = 0.224, p < 0.001; Maximum Parsimony phylogeny: Blomberg’s *K* = 0.649, p = 0.05), as well for T_max_ (Bayesian phylogeny: Blomberg’s *K* = 0.833, p < 0.001; Maximum likelihood phylogeny: Blomberg’s *K* = 0.267, p < 0.001; Maximum Parsimony phylogeny: Blomberg’s *K* = 1.87, p < 0.001). While closely related *Phytophthora* species had more similar thermal physiology than random pairs, in all but one case differences were greater than that expected under a Brownian motion evolutionary model (*27, 28*). This could not be explained by known geographical locations of species (Fig S1). Apparent latitudinal range shifts of plant pathogens in response to global warming (*29*) suggest niche conservatism in thermal physiology, i.e. migration is the dominant response of populations to changing climates rather than adaptation to new climates in situ (*30*). There is limited evidence for thermal adaptation in fungal pathogens (*31*) and within *Phytophthora* (*32*). Our analysis suggests limited phylogenetic constraint in temperature niche evolution in *Phytophthora*, though we acknowledge that a significant but small phylogenetic signal (*K* < 1) can arise from several different evolutionary models (*33*).

In the evolution of host range, closely-related plant species share pathogens (*34*) but the degree to which closely-related pathogens share plant hosts is unclear. *Formae speciales* of powdery mildews, for example, are specialized upon, but not restricted to, particular plant hosts (*35*). Evidence for different types of cophylogenetic dynamics in plant-fungus symbioses ranging from close congruence indicating codivergence to incongruence indicating long-range host switching (*36*). Host jumps (acquisition of a host phylogenetically distant from current hosts) and transitions from specialist to generalist or vice-versa are known in plant pathogens (*37, 38*), suggesting that a host range evolution could be more evolutionarily labile than temperature physiology. We found a statistically significant cophylogenetic association between the topologies of three *Phytophthora* phylogenies and the phylogeny of their plant hosts (436 species-level pathogen-host interaction records; Bayesian phylogeny, m^2^_XY_ = 0.952, p < 0.001; Maximum likelihood phylogeny: m^2^_XY_ = 0.951, p < 0.001; Maximum Parsimony phylogeny: m^2^_XY_ = 0.946, p < 0.001) (Fig. S2). Wide-ranging host jumps _are known_ in *Phytophthora*, for example clade 1c (*P. infestans, P. ipomoeae, P. mirabilis*, and *P. phaseoli*) evolved through an ancestral major host jump, followed by adaptive specialisation to one of four plant families respectively (*39*). *P. infestans* has been recorded on 22 hosts, 20 in the Solanaceae and 2 in the sister-family Convolvulaceae, while *P. cactorum* infects the gymnosperm *Abies balsamea* as well as diverse angiosperms.

Our analysis of fungal and oomycete plant-associated cardinal temperatures shows that abiotic Fundamental Niches are both wider and different in form than corresponding Realized Niches. We propose that this demonstrates the expected restriction in the breadth of the Realized Niche in comparison to the Fundamental Niche (*1*). We show that microbial specialization can occur independently in biotic and abiotic niche axes, suggesting that the terms “specialist” and “generalist” should be used cautiously when describing the ecology of microbial species. Finally, we show that that both the thermal niche and host ranges are evolutionarily labile in genus *Phytophthora*, but retain phylogenetic signal. Invasive fungi and oomycetes are spreading rapidly around the world to challenge global food security, partly in response to climate change (*29*). Therefore, the shape and size of the microbial niche has important implications for the management of natural and agricultural ecosystems.

## References

1. J. M. Chase, M. A. Leibold, Ecological Niches: Linking Classical and Contemporary Approaches (University of Chicago Press, Chicago, 2003; https://www.press.uchicago.edu/ucp/books/book/chicago/E/bo3638660.html).

2. G. E. Hutchinson, Concluding Remarks. Cold Spring Harb Symp Quant Biol. 22, 415–427 (1957).

3. R. K. Colwell, T. F. Rangel, Hutchinson’s duality: The once and future niche. PNAS. 106, 19651–19658 (2009).

4. B. Blonder, C. Lamanna, C. Violle, B. J. Enquist, The n-dimensional hypervolume. Global Ecology and Biogeography. 23, 595–609 (2014).

5. J. Elith, M. Kearney, S. Phillips, The art of modelling range-shifting species. Methods in Ecology and Evolution. 1, 330–342 (2010).

6. J. Soberón, M. Nakamura, Niches and distributional areas: concepts, methods, and assumptions. Proceedings of the National Academy of Sciences. 106, 19644–19650 (2009).

7. J. P. DeLong, J. P. Gibert, T. M. Luhring, G. Bachman, B. Reed, A. Neyer, K. L. Montooth, The combined effects of reactant kinetics and enzyme stability explain the temperature dependence of metabolic rates. Ecology and Evolution. 7, 3940–3950 (2017).

8. P. M. Schulte, The effects of temperature on aerobic metabolism: towards a mechanistic understanding of the responses of ectotherms to a changing environment. Journal of Experimental Biology. 218, 1856–1866 (2015).

9. M. B. Araújo, M. Luoto, The importance of biotic interactions for modelling species distributions under climate change. Global Ecology and Biogeography. 16, 743–753 (2007).

10. R. P. Anderson, When and how should biotic interactions be considered in models of species niches and distributions? Journal of Biogeography. 44, 8–17 (2017).

11. R. Early, S. A. Keith, Geographically variable biotic interactions and implications for species ranges. Global Ecology and Biogeography. 28, 42–53 (2019).

12. J. P. Sexton, J. Montiel, J. E. Shay, M. R. Stephens, R. A. Slatyer, Evolution of Ecological Niche Breadth. Annu. Rev. Ecol. Evol. Syst. 48, 183–206 (2017).

13. C. B. de Araújo, L. O. Marcondes-Machado, G. C. Costa, The importance of biotic interactions in species distribution models: a test of the Eltonian noise hypothesis using parrots. Journal of Biogeography. 41, 513–523 (2014).

14. H. R. Cunningham, L. J. Rissler, L. B. Buckley, M. C. Urban, Abiotic and biotic constraints across reptile and amphibian ranges. Ecography. 39, 1–8 (2016).

15. C. González-Salazar, C. R. Stephens, P. A. Marquet, Comparing the relative contributions of biotic and abiotic factors as mediators of species’ distributions. Ecological Modelling. 248, 57–70 (2013).

16. L. R. Thompson, et al., A communal catalogue reveals Earth’s multiscale microbial diversity. Nature. 551, 457 (2017).

17. M. Delgado-Baquerizo, F. T. Maestre, P. B. Reich, T. C. Jeffries, J. J. Gaitan, D. Encinar, M. Berdugo, C. D. Campbell, B. K. Singh, Microbial diversity drives multifunctionality in terrestrial ecosystems. Nature Communications. 7, 10541 (2016).

18. S. Barton, G. Yvon-Durocher, Quantifying the temperature dependence of growth rate in marine phytoplankton within and across species. Limnology and Oceanography. 0, doi:10.1002/lno.11170.

19. L. A. Stevenson, R. A. Alford, S. C. Bell, E. A. Roznik, L. Berger, D. A. Pike, Variation in Thermal Performance of a Widespread Pathogen, the Amphibian Chytrid Fungus Batrachochytrium dendrobatidis. PLOS ONE. 8, e73830 (2013).

20. C. J. Alster, Z. D. Weller, J. C. von Fischer, A meta-analysis of temperature sensitivity as a microbial trait. Global Change Biology. 24, 4211–4224 (2018).

21. A. I. Dell, S. Pawar, V. M. Savage, The thermal dependence of biological traits. Ecology. 94, 1205–1206 (2013).

22. D. Storch, L. Menzel, S. Frickenhaus, H.-O. Pörtner, Climate sensitivity across marine domains of life: limits to evolutionary adaptation shape species interactions. Global Change Biology. 20, 3059–3067 (2014).

23. W. Yan, L. A. Hunt, An Equation for Modelling the Temperature Response of Plants using only the Cardinal Temperatures. Ann Bot. 84, 607–614 (1999).

24. A.-L. Boixel, G. Delestre, J. Legeay, M. Chelle, F. Suffert, Phenotyping Thermal Responses of Yeasts and Yeast-like Microorganisms at the Individual and Population Levels: Proof-of-Concept, Development and Application of an Experimental Framework to a Plant Pathogen. Microb Ecol. 78, 42–56 (2019).

25. E. J. De Jong, A. B. Eskes, J. G. J. Hoogstraten, J. C. Zadoks, Temperature requirements for germination, germ tube growth and appressorium formation of urediospores of Hemileia vastatrix. Netherlands Journal of Plant Pathology. 93, 61–71 (1987).

26. X. Yang, B. M. Tyler, C. Hong, An expanded phylogeny for the genus Phytophthora. IMA Fungus. 8, 355–384 (2017).

27. D. Ackerly, Conservatism and diversification of plant functional traits: Evolutionary rates versus phylogenetic signal. PNAS. 106, 19699–19706 (2009).

28. S. P. Blomberg, T. Garland, Tempo and mode in evolution: phylogenetic inertia, adaptation and comparative methods. Journal of Evolutionary Biology. 15, 899–910 (2002).

29. D. P. Bebber, Range-Expanding Pests and Pathogens in a Warming World. Annu. Rev. Phytopathol. 53, 335–356 (2015).

30. D. Nogués-Bravo, F. Rodríguez-Sánchez, L. Orsini, E. de Boer, R. Jansson, H. Morlon, D. A. Fordham, S. T. Jackson, Cracking the Code of Biodiversity Responses to Past Climate Change. Trends in Ecology & Evolution. 33, 765–776 (2018).

31. J. Zhan, B. A. McDonald, Thermal adaptation in the fungal pathogen Mycosphaerella graminicola. Molecular Ecology. 20, 1689–1701 (2011).

32. X. Yang, M. E. Gallegly, C. Hong, A high-temperature tolerant species in clade 9 of the genus Phytophthora: P. hydrogena sp. nov. Mycologia. 106, 57–65 (2014).

33. L. J. Revell, L. J. Harmon, D. C. Collar, Phylogenetic Signal, Evolutionary Process, and Rate. Syst Biol. 57, 591–601 (2008).

34. G. S. Gilbert, I. M. Parker, The Evolutionary Ecology of Plant Disease: A Phylogenetic Perspective. Annual Review of Phytopathology. 54, 549–578 (2016).

35. V. Troch, K. Audenaert, R. A. Wyand, G. Haesaert, M. Höfte, J. K. M. Brown, Formae speciales of cereal powdery mildew: close or distant relatives? Molecular Plant Pathology. 15, 304–314 (2014).

36. A. P. Jackson, A Reconciliation Analysis of Host Switching in Plant-Fungal Symbioses. Evolution. 58, 1909–1923 (2004).

37. L. G. Barrett, M. Heil, Unifying concepts and mechanisms in the specificity of plant– enemy interactions. Trends in Plant Science. 17, 282–292 (2012).

38. C. E. Morris, B. Moury, Annual Review of Phytopathology, in press, doi:10.1146/annurev-phyto-082718-100034.

39. S. Raffaele, R. A. Farrer, L. M. Cano, D. J. Studholme, D. MacLean, M. Thines, R. H. Y. Jiang, M. C. Zody, S. G. Kunjeti, N. M. Donofrio, B. C. Meyers, C. Nusbaum, S. Kamoun, Genome Evolution Following Host Jumps in the Irish Potato Famine Pathogen Lineage. Science. 330, 1540–1543 (2010).

