## Supplementary Material for "Geometry and evolution of the ecological niche in plant-associated microbes"

### Materials and Methods

Cardinal temperature data collection

Experimentally-derived minimum ($T_{min}$), optimum ($T_{opt}$) and maximum ($T_{max}$) temperatures (collectively 'cardinal temperatures') of five life cycle processes (disease development (DD), fructification (FR), infection (IN), spore germination (SG) and sporulation (SP)), as well as axenic growth in culture (GC) (collectively 'biological processes'), were extracted from ref. (*39*) for fungi and oomycete species, hereafter referred to as the 'Togashi dataset' (Data S1). Additionally, GC $T_{opt}$ and $T_{max}$ data were extracted for 104 *Phytophthora* species from (*40*), hereafter referred to as the 'Martin dataset'. Finally, IN cardinal temperature for 45 plant pathogens were extracted from ref. (*41*), hereafter referred to as the 'Magarey dataset'.

The Species Fungorum database (SFD) (www.speciesfungorum.org) was used to identify synonymous names for each species recorded in the Togashi dataset [accessed between 1/5/2017 – 18/10/2019]. Where no information was available on the SFD, or discovery and/or sanction author(s) (where available) were greatly inconsistent, Mycobank (www.mycobank.org) was used as an alternative [accessed between 1/5/2017 – 18/10/2019]. If a species could not be identified on the SFD or Mycobank it was excluded from the dataset. Discovery and sanction author(s) of species were used (where available) to improve accuracy of matching synonymous species names between those recorded in (*39*) and the SFD/Mycobank. Species were recorded by their current name, according to either SFD or Mycobank. Where spelling of species names differed between ref. (*39*) and the SFD/Mycobank, but it was likely that the spelling in ref. (*39*) was an error, species names were updated to reflect this. In some cases, species were recorded in ref. (*39*) under multiple, synonymous names. However, said species names were at times found to be not synonymous, once identified in the SFD and/or Mycobank. Various methods were used to correct for this, detailed below.

If cardinal temperature data associated with multiple, nonsynonymous species did not specify which species the data referred to, data were recorded under the first species given in ref. (*39*). For example, in ref. (*39*) GC and DD cardinal temperature data were recorded for *Mycosphaerella tulasnei* (Jancz.) [syn. *Cladosporium herbarum* (Link)]. The SFD classified *Mycosphaerella tulasnei* (Jancz.) Lindau (1903) as *Mycosphaerella tassiana* (De Not.) Johanson (1884), but *Cladosporium herbarum* (Pers.) Link (1816) as *Cladosporium herbarum* (Pers.) Link (1816). However, it was not clear in ref. (*39*) which cardinal temperatures referred specifically to which species. Hence, all data were recorded under *Mycosphaerella tassiana*.

In contrast, where cardinal temperature data specifically referred to a nonsynonymous species, these were recorded under the correct species name. For example, GC cardinal temperature data were recorded for *Fusarium solani* var. *martii* (Appr. et Wr.) Wr. [syn.

*Fusarium martii* var. *phaseoli* (Buskh)]. The SFD classified *Fusarium solani* var. *martii* (Appel & Wollenw.) Wollenw. (1930) as *Fusarium solani* (Mart.) Sacc. (1881), but *Fusarium martii* var. *phaseoli* (Burkh. 1919) as *Neocosmospora phaseoli* (Burkh.) L. Lombard & Crous 2015. However, a subset of cardinal temperature data was recorded as "as *Mart. Phaseoli*" and so was assigned to *Neocosmospora phaseoli*, and not *Fusarium solani*.

Finally, all cardinal temperature data concerning *Fusarium oxysporum* formae speciales were recorded under *Fusarium oxysporum*, as well as their respective formae speciales. *Formae speciales* were excluded from analyses concerning the shape of the abiotic niche (see below), but included in the analysis of niche co-specialisation, thereby maintaining strain-host interactions in this analysis.

The methods of determining species names detailed above resulted in cardinal temperature data for 661 species (599 fungi and 62 oomycetes, n = 8231) being recorded in the Togashi dataset. All information regarding how species were named in the Togashi dataset, including discovery name author(s)/sanction name author(s), changes to spelling of species names, synonymous and nonsynonymous species names, and cases where data were extracted from one species record and separately recorded as a different species can be found in the Data S1 (*38*).

For each data point recorded in the Togashi dataset, where ref. (*39*) recorded that the 'true' value lies above or below the value provided, the value provided was recorded. For example, if $T_{min}$ was recorded as 'below 8 °C', 8 °C was recorded as $T_{min}$; if $T_{max}$ was recorded as 'above 25 °C', 25 °C was recorded as $T_{max}$. Where a range was provided, the mid-point was recorded. However, where a range was provided, but the 'true' value was recorded to lie above or below this, the upper or lower limit was chosen, respectively. For example, if $T_{opt}$ was quoted as 'below 18 - 20 °C', 18 °C was recorded as $T_{opt}$. Where a range was quoted for the entire biological process, the upper and lower bounds were recorded as $T_{max}$ and $T_{min}$, respectively. For example, if IN was quoted as 'occurring between 5 - 35 °C', 5 °C was recorded as $T_{min}$ and 35 °C was recorded as $T_{max}$, except where it was apparent that the temperature range quoted referred to optimal conditions. In such cases, the mid-point was recorded as $T_{opt}$. Data regarding 'Infection and disease development' were independently included in the dataset under IN and DD, unless it was apparent that the data likely related to specifically to IN or DD. Data quoted in (*39*) that was the result of complex treatments and/or was not obviously related to $T_{min}$, $T_{opt}$, or $T_{max}$ were excluded. Further information regarding how each data point in the Togashi dataset was determined can be found in Data S1 (*38*). Where multiple references were provided for a single data point in ref. (*39*), this was taken to represent independent observations, and so were individually included in the Togashi dataset. Species taxonomic classification was determined using either the SFD or Mycobank [accessed between 1/5/2017 - 18/10/2019]. All data extraction was completed by the same researcher. For the Magarey dataset, pathogen names were updated according to the SFD or Mycobank [accessed between 1/5/2017 - 18/10/2019] to ensure correct matching to the Togashi dataset. No additional processing was performed on the Martin dataset.

Data analysis

All analyses were performed in R 3.2.3 (*42*). In all analyses the mean of $T_{min}$, $T_{opt}$ or $T_{max}$ for a given biological process, for a given species, is treated as a single datapoint.

Data Validation

15 pathogens were recorded in both the Togashi and Magarey datasets. For these species, root mean square error (RMSE) was calculated between IN cardinal temperature estimates. When all data was included, RMSE was calculated as 5.36 °C (n = 41) (Fig. S3a). Clustering of data points at 35 °C and 1 °C along the y-axis is a result of how ref. (*41*) estimated $T_{max}$ and $T_{min}$, respectively - if no $T_{max}$ for infection was found, the authors set $T_{max}$ to 35 °C. Similarly, if no $T_{min}$ for infection was found, but infection could occur lower than the hosts developmental threshold, the authors set $T_{min}$ to be 5 °C lower than the lowest tested temperature, but not lower than 1 °C. When bounded $T_{min}$ and $T_{max}$ data was excluded, RMSE was calculated as 5.07 °C (n = 27) (Fig. S3b). Fig. S3a,b indicate that the greatest deviation from an identity relationship (dotted line) arises around $T_{min}$. This is likely due to $T_{min}$ being more problematic to quantify - the lowest temperature a given biological process occurs at will depend on the amount of time given for the process to occur, whereas $T_{max}$ is likely to be clearly defined as enzymes denature. Further, 22 *Phytophthora* species were present in both the Togashi and Martin datasets. For these pathogens, RMSE was calculated for GC $T_{opt}$ as 2.51 °C (n = 20) (Fig. S3c) and GC $T_{max}$ as 3.42 °C (n = 22) (Fig. S3d).

<u>Analysis of species cardinal temperature</u>

The Togashi dataset was used for this analysis. $T_{range}$ was calculated as the range between $T_{min}$ and $T_{max}$. $T_{range(50\%)}$ was calculated as the range between $T_{min(50\%)}$ and $T_{max(50\%)}$; $T_{max(50\%)}$ and $T_{min(50\%)}$ refer to $T_{max}$ and $T_{min}$ where a species response rate = 0.5 (at $T_{opt}$ the responses = 1, at $T_{min}$ and $T_{max}$ the response = 0). Hence, $T_{range(50\%)}$ reflects the temperature range where a species performs a biological process well. Responses were calculated by a beta function (Equation S1) that uses a species' cardinal temperature to estimate a temperature performance curve (*43*). In some cases, for particular species-biological process combinations, mean $T_{opt}$ was estimated as greater than mean $T_{max}$ or lower than mean $T_{min}$. This is because data were extracted from multiple sources. $T_{range}$, $T_{range(50\%)}$, and skew were not calculated for these species-biological process combinations.

$$\text{Equation S1:} \, r(\mathrm{T}) = \left(\frac{\mathrm{T}_{max} - \mathrm{T}}{\mathrm{T}_{max} - \mathrm{T}_{opt}}\right)\left(\frac{\mathrm{T} - \mathrm{T}_{min}}{\mathrm{T}_{opt} - \mathrm{T}_{min}}\right)^{(\mathrm{T}_{opt}-\mathrm{T}_{min})/(\mathrm{T}_{max}-\mathrm{T}_{opt})}$$

Species with at least one $T_{opt}$, $T_{range}$ or skew estimate were included in analyses involving $T_{opt}$, $T_{range}$ and skew, respectively. Skew (s) was calculated according to Equation S1. Where skew > 0.5, $T_{opt}$ is closer to $T_{max}$ than $T_{min}$; where skew < 0.5, $T_{opt}$ is closer to $T_{min}$ than $T_{max}$. $T_{min}$, $T_{opt}$, $T_{max}$, and $T_{range}$ were analysed individually by linear mixed effects analysis (*44*). Biological process and species classification (fungi or oomycete) were entered as fixed effects and species name was entered a random effect. Sample size varies between box plots and scatter plots as a species may have a $T_{min}$, $T_{opt}$ and/or $T_{max}$ estimate for one biological process, but not others.

$$\text{Equation S2:} \, \mathrm{s} = \frac{\mathrm{T}_{opt} - \mathrm{T}_{min}}{\mathrm{T}_{max} - \mathrm{T}_{min}}$$

<u>Analysis of abiotic and biotic niche co-specialisation</u>

The Togashi dataset was used for this analysis. The PlantWise database (CABI) [accessed 28/10/2013, by permission] provides information on known pathogen/host interactions. To ensure correct pathogen species matching between the Togashi dataset and the PlantWise database, some pathogen species names were updated in PlantWise database (Table S6), according to their respective, current names given in the SFD and/or Mycobank. 236 species

(198 fungi, 38 oomycetes) were identified in the PlantWise database with at least one $T_{range}$ ($T_{range(50\%)}$) estimate, for at least one biological process in the Togashi dataset. Plant hosts were identified to species level i.e. varieties were excluded. Two different methods were used to quantify pathogen host diversity. First, any hosts not recorded in the PlantWise database to species level were excluded. In this case, 983 hosts of 231 pathogens were utilised to generate a time-calibrated host phylogeny using the R function 'S.PhyloMaker' (scenario 1, genera or species are added as basal polytomies within their families or genera) (*45*). The resultant pathogen/host database and generated host phylogeny are hereafter referred to as the 'unprocessed pathogen/host database' and 'unprocessed host phylogeny' (Fig. S4) respectively. Second, where a host record in the PlantWise database was not identified to species level, it was assumed that the pathogen in question was able to successfully infect all species identified in S.PhyloMaker within that taxonomic rank. For example, *Leptosphaeria maculans* was recorded in the PlantWise database as being a pathogen of the order Gentianales. Hence, 1388 host species found within the order Gentianales in S.PhyloMaker were added to *L. maculans* host range. The resultant pathogen/host database and generated host phylogeny are referred to as the 'processed pathogen/host database' and 'processed host phylogeny' respectively. In this case, 11,983 hosts of 235 pathogens were used to generate a time-calibrated host phylogeny, also using the R function 'S.PhyloMaker' (scenario 1) (*45*). For both methods, to ensure correct matching of host names between the pathogen/host databases and S.PhyloMaker, some corrections to host species names in the PlantWise database were made (Table S7). Eight hosts were not identifiable in S.PhyloMaker during phylogeny construction, and were excluded from the analysis (Table S7). This was due to either uncertainty in classification and phylogenetic position, or due to missing data in S.PhyloMaker. The function 'pd' in the R package 'picante' (*46*) was used to quantify host diversity of each pathogen. Host diversity was calculated as Faith's phylogenetic diversity (PD) (*47*). The phylogeny root node was excluded in all calculations. Hence, pathogens with only one host scored a PD of 0. Co-specialistion across abiotic ($T_{range}$ or $T_{range(50\%)}$) and biotic ($\log_{10+1}$-transformed host diversity) niche axes was calculated by Pearson correlation.

<u>Analysis of *Phytophthora* species cardinal temperature phylogenetic signal</u>

The Martin dataset was used for this analysis. Phylogenies constructed by (1) Bayesian, (2) Maximum likelihood and (3) Maximum Parsimony methods for *Phytophthora* species were extracted from (*25*) (TreeBASE S19303). 99 *Phytophthora* species (*P. alticola*, *P. andina*, *P. aquimorbida*, *P. arenaria*, *P. austrocedrae*, *P. bisheria*, *P. boehmeriae*, *P. botryose*, *P. brassicae*, *P. cactorum*, *P. cajani*, *P. cambivora*, *P. capensis*, *P. capsici*, *P. captiosa*, *P. chrysanthemi*, *P. cinnamomi*, *P. citricola*, *P. citrophthora*, *P. clandestina*, *P. colocasiae*, *P. constricta*, *P. cryptogea*, *P. drechsleri*, *P. elongata*, *P. erythroseptica*, *P. europaea*, *P. fallax*, *P. fluvialis*, *P. foliorum*, *P. fragariae*, *P. frigida*, *P. gallica*, *P. gemini*, *P. gibbosa*, *P. glovera*, *P. gonapodyides*, *P. gregata*, *P. hedraiandra*, *P. heveae*, *P. hibernalis*, *P. humicola*, *P. hydropathica*, *P. idaei*, *P. ilicis*, *P. infestans*, *P. inflata*, *P. insolita*, *P. inundata*, *P. ipomoea*, *P. iranica*, *P. irrigata*, *P. kernoviae*, *P. lateralis*, *P. litoralis*, *P. macrochlamydospora*, *P. meadii*, *P. medicaginis*, *P. megakarya*, *P. megasperma*, *P. melonis*, *P. mengei*, *P. mexicana*, *P. mirabilis*, *P. morindae*, *P. multivesiculata*, *P. multivora*, *P. nemorosa*, *P. nicotianae*, *P. obscura*, *P. palmivora*, *P. parsiana*, *P. phaseoli*, *P. pini*, *P. pinifolia*, *P. pistaciae*, *P. plurivora*, *P. polonica*, *P. primulae*, *P. pseudosyringae*, *P. pseudotsugae*, *P. psychrophila*, *P. quercetorum*, *P. quercina*, *P. quininea*, *P. ramorum*, *P. richardiae*, *P. rosacearum*, *P. rubi*, *P. sansomeana*, *P. siskiyouensis*, *P. sojae*, *P. syringae*, *P. tentaculata*, *P. thermophila*, *P. trifolii*, *P. tropicalis*, *P. uliginosa*, and *P. vignae*) were present in both the Martin dataset and extracted phylogenies. The function 'phylosig' in the R package 'phytools' (*48*) was used to separately test for a phylogenetic signal for GC $T_{opt}$ and $T_{max}$. 10,000 simulations were assigned in each analysis for randomization test. Where multiple strains of a particular species were included in a phylogeny, only one strain was assigned a GC $T_{opt}$ or $T_{max}$ record from the Martin dataset, thereby preventing pseudoreplication.

The influence of spatial autocorrelation on phylogenetic signal of *Phytophthora* species cardinal temperature was also investigated. For 30 *Phytophthora* species included in the above analysis (*P. boehmeriae*, *P. botryosa*, *P. cactorum*, *P. cambivora*, *P. capsici*, *P. cinnamomi*, *P. citrophthora*, *P. colocasiae*, *P. cryptogea*, *P. drechsleri*, *P. erythroseptica*, *P. fragariae*, *P. infestans*, *P. kernoviae*, *P. lateralis*, *P. macrochlamydospora*, *P. meadii*, *P. medicaginis*, *P. megakarya*, *P. megasperma*, *P. nicotianae*, *P. palmivora*, *P. pseudosyringae*, *P. quercetorum*, *P. quercina*, *P. ramorum*, *P. rubi*, *P. sojae*, and *P. vignae*), estimates of presence at country or region scale were extracted from the CABI Distribution Maps of Plant Diseases (*49, 50*). The centroid of country or region were used for all records. Mantel correlations (MCs) were performed between GC $T_{opt}$ (and $T_{max}$) distance and great circle distance (km) or average air surface temperature (AST) distance (°C) matrices. Average AST was extracted from Berkley Earth (http://www.berkeleyearth.org) for each latitude-longitude location. Latitude and longitude from ref (*49, 50*) were rounded to the nearest 0.5 ° to align with those extracted from Berkley Earth for analysis of average air surface temperature distance. All MC were performed using the function 'mantel' in the R package 'ecodist' (*51*) with 10,000 iterations to calculate bootstrapped confidence limits.

Analysis of cophylogenetic association between *Phytophthora* and plant host species

The Martin dataset was used for this analysis. 34 *Phytophthora* species were present in both the PlantWise database and extracted *Phytophthora* phylogenies detailed above (*P. alni*, *P. asparagi*, *P. boehmeriae*, *P. botryose*, *P. cactorum*, *P. cambivora*, *P. capsica*, *P. cinnamomi*, *P. citricola*, *P. citrophthora*, *P. colocasiae*, *P. cryptogea*, *P. drechsleri*, *P. erythroseptica*, *P. fragariae*, *P. hibernalis*, *P. infestans*, *P. kernoviae*, *P. lateralis*, *P. macrochlamydospora*, *P. meadii*, *P. medicaginis*, *P. megakarya*, *P. megasperma*, *P. nicotianae*, *P. palmivora*, *P. phaseoli*, *P. pseudotsugae*, *P. ramorum*, *P. richardiae*, *P. rubi*, *P. sojae*, *P. syringae*, and *P. vignae*). 256 hosts recorded to species level in the PlantWise database were extracted (i.e. those recorded to genus were excluded and host variety was ignored) and utilised to generate a time-calibrated host phylogeny using the R function 'S.PhyloMaker' (scenario 1) (*45*). The function 'PACo' in the R package 'paco' (*52*) was used to test for cophylogenetic association between each *Phytophthora* species phylogeny and the time-calibrated host phylogeny. The method 'quasiswap' was assigned, which is a more constrained method than others available, where the number of interactions is conserved for each species (and hence in the network as a whole). This method was used because we make no assumption about which group (host or pathogen) is tracking the other (*53*). 10,000 randomisations were assigned in each analysis. Where multiple strains of a particular species were included in a phylogeny, only one strain was assigned a host range, thereby preventing pseudoreplication.

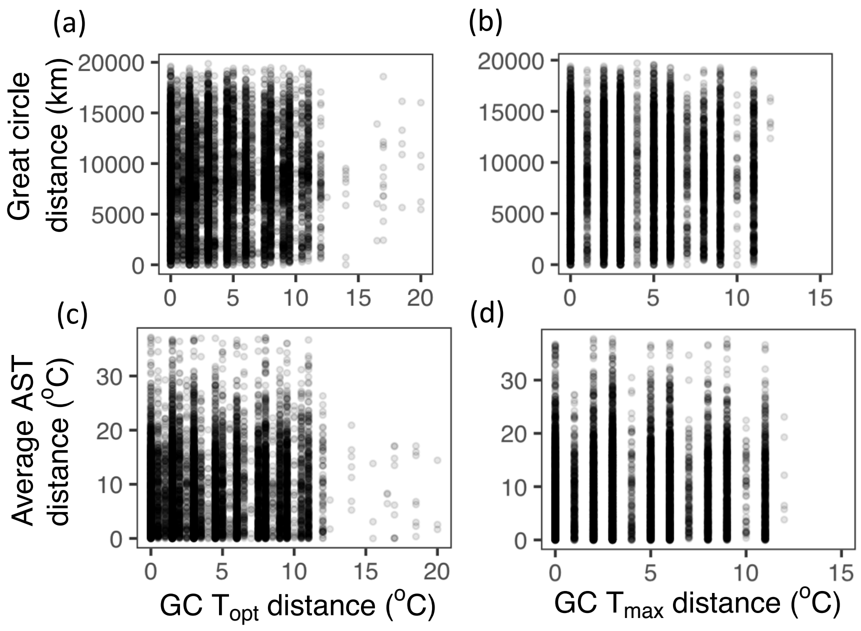


**Fig. S1.**

Analysis of spatial correlation on phylogenetic signal of *Phytophthora* species cardinal temperatures. (a, c) GC $T_{opt}$. (b, d) GC $T_{max}$. (b, d) Relationship between GC temperature response distance (°C) and great circle distance (km). (a) Mantel correlation (MC) = -0.017, $p < 0.01$. (b) MC = 0.011, $p < 0.05$. (c, d) Relationship between GC temperature response distance (°C) and average air surface temperature (AST) (°C). (c) MC = 0.020, $p < 0.05$. (d) MC = 0.042, $p = 0.001$. Graphs show 10,000 randomly sampled datapoints. $n_{species} = 29$; $n_{distances} = 1,319,500$.

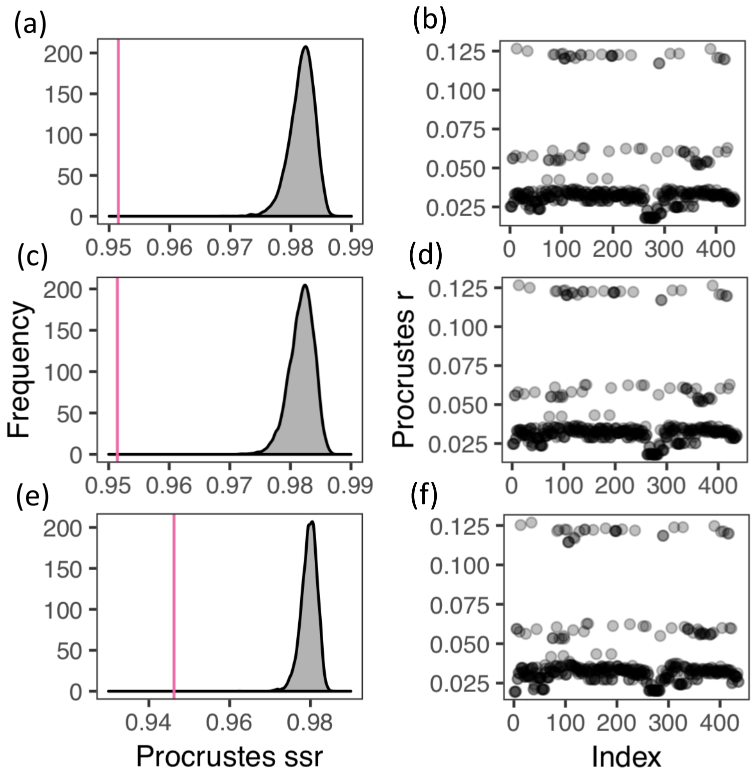


**Fig. S2.**

Cophylogenetic association analysis outputs. (a, c, e) The observed best-fit Procrustean super-imposition (pink lines) (0.952, 0.951, 0.946) were better than the same for any of the ensemble of network randomisations in each null model (n = 10,000). (b, d, f) Procrustes residuals for all interactions in each pathogen-host network (n = 436). (a, b) Bayesian *Phytophthora* spp. phylogeny. (c, d) Maximum likelihood *Phytophthora* spp. phylogeny. (e, f) Maximum parsimony *Phytophthora* spp. phylogeny. All *Phytophthora* spp. phylogenies were extracted form (*25*) (TreeBASE S19303).

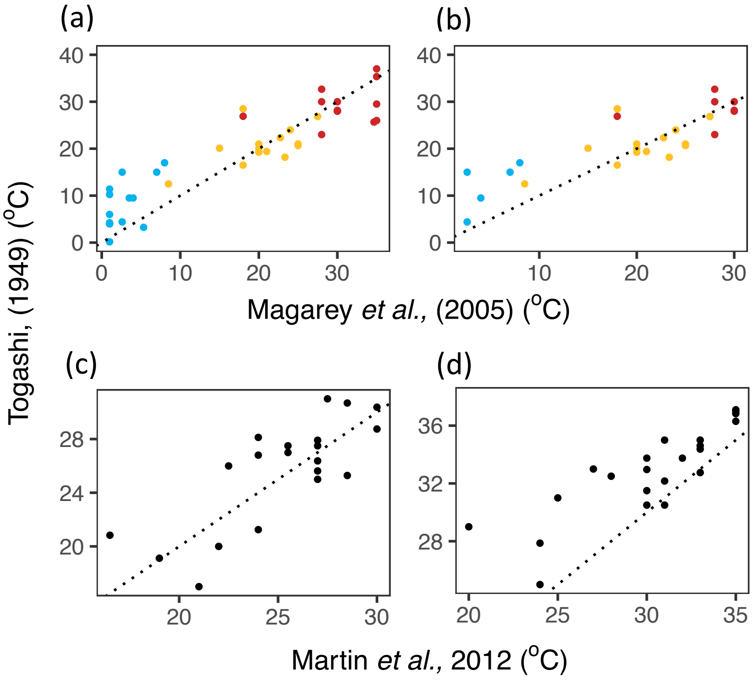


**Fig. S3.**

Relationship between infection CT estimates from (*39*) and (*41*). (a) Inclusion of all available data, root mean square error (RMSE) = 5.36 °C, n = 41. (b) Exclusion of bounded Tmin and Tmax (1 °C and 35 °C) data in (41), RMSE = 5.07 °C, n = 27. (a, b) Colour refers to estimates of $T_{min}$ (blue), $T_{opt}$ (yellow) and $T_{max}$ (red). (c) Relationship between GC $T_{opt}$ and (d) $T_{max}$ estimates from (*39*) and (*40*). (c) RMSE = 2.51 °C, n = 20. (d) RMSE = 3.42 °C, n = 22. Dotted line indicates identity relationship.

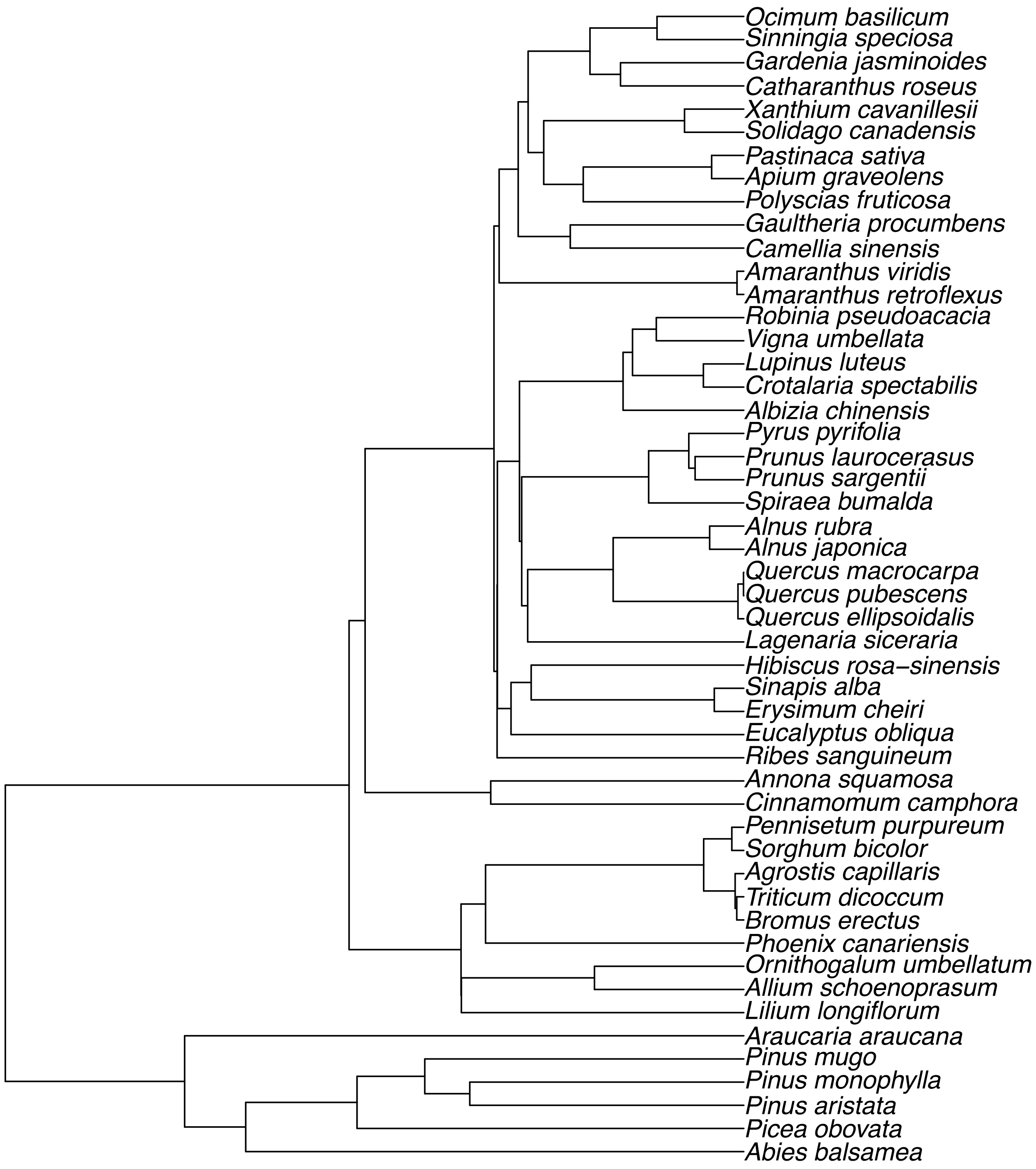


**Fig. S4.**

Time-calibrated unprocessed host phylogeny constructed using S. PhyloMaker. 50 tips randomly extracted (total tips in constructed phylogeny = 983). Only hosts recorded to species level in the PlantWise database are included.

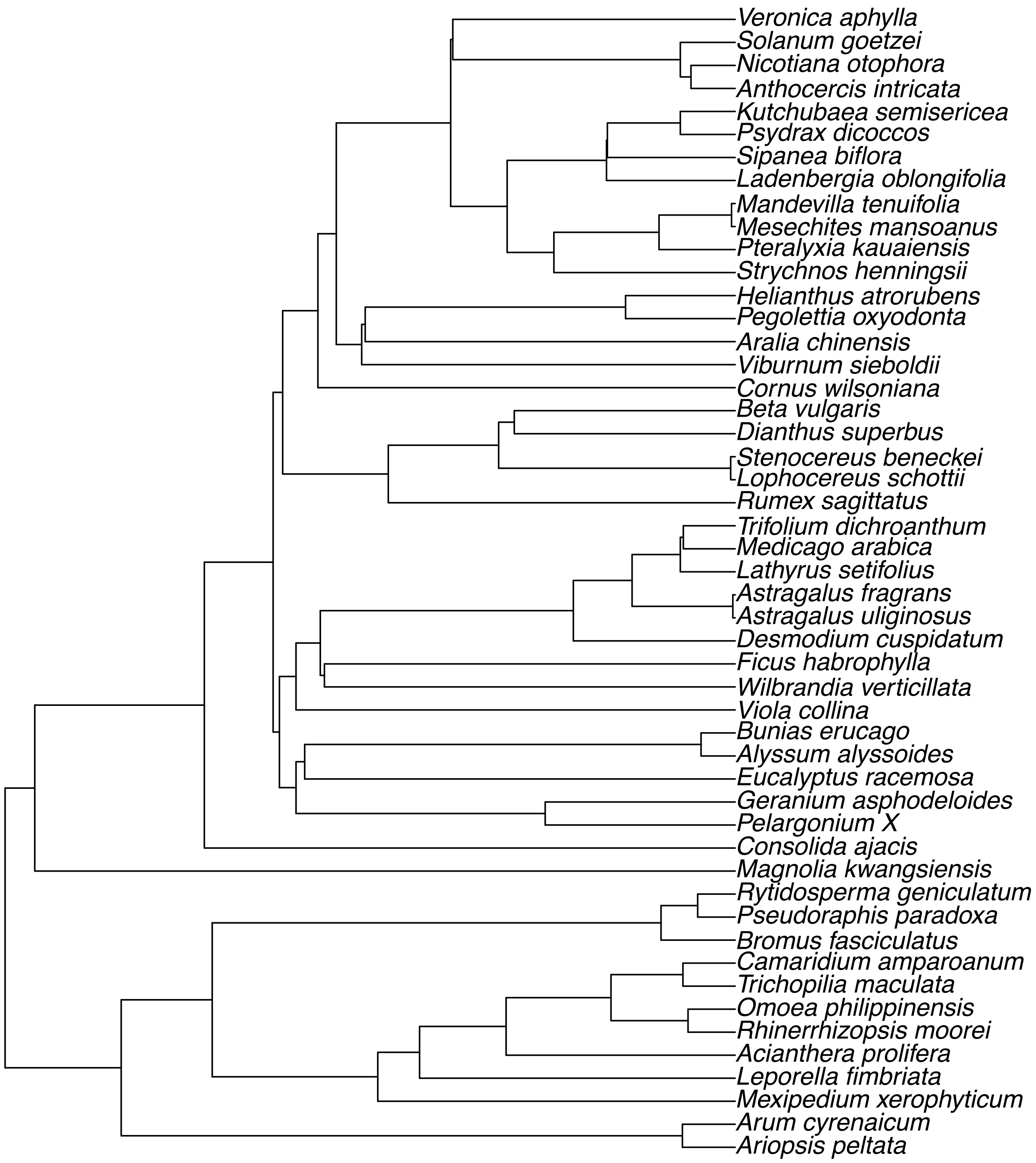


**Fig. S5.**

Time-calibrated processed host phylogeny constructed using S. PhyloMaker. 50 tips randomly extracted (total tips in constructed phylogeny = 15,460). Only hosts recorded to species level in the PlantWise database are included.

**Table S1.**

Summary of temperature responses of fungi and oomycetes for different processes. Medians with interquartile ranges in parentheses.

| | Fungi | | |
|---|---|---|---|
| Process | $T_{min}$ | $T_{opt}$ | $T_{max}$ |
| GC | 6.0 (4.0-9.4) | 25.5 (22.6 – 28.0) | 35 (32.5 – 37.5) |
| DD | 10.0 (4.9 – 14.0) | 22.3 (19.5 – 26.3) | 30.4 (27.2 – 32.8) |
| FR | 10.0 (7.3 – 12.1) | 21.7 (17.7 – 25.0) | 30.0 (27.0 – 31.25) |
| IN | 10.0 (7.0 – 15.0) | 21.0 (18.1 – 26.9) | 30.0 (26.7 – 34.0) |
| SG | 5.8 (3.0 – 8.1) | 23.0 (19.0 – 26.0) | 32.2 (30.0 – 36.0) |
| SP | 11.5 (4.8 – 15.0) | 23.8 (19.1 – 27.2) | 31.5 (27.0 – 34.2) |

| | Oomycetes | | |
|---|---|---|---|
| Process | $T_{min}$ | $T_{opt}$ | $T_{max}$ |
| GC | 8.0 (6.3 – 9.0) | 27.0 (25.8 – 0.0) | 34.8 (32.8 – 37.0) |
| DD | 9.9 (5.7 – 12.4) | 22.4 (20.1 – 26.0) | 30.0 (29.5 – 32.3) |
| FR | 10.9 (7.9 -13.8) | 21.5 (17.4 – 24.3) | 25.7 (21.2 – 32.9) |
| IN | 10.3 (4.4 – 13.8) | 20.6 (19.8 – 23.3) | 31.5 (29.8 – 34.0) |
| SG | 6.5 (4.0 – 9.9) | 19.0 (15.9 – 23.8) | 28.8 (26.1 – 31.8) |
| SP | 7.4 (5.5 - 8.7) | 18.3 (15.6 – 19.4) | 28.8 (23.9 – 29.5) |

**Table S2.**
Difference between temperature responses for growth in culture (GC) and other processes within species. Positive values indicate that the alternative process has a higher value than GC. Fungi and oomycetes are not differentiated because only a small number of pairwise comparisons were available for oomycetes.

| Response | Process | Difference (°C) | df | t | p |
|---|---|---|---|---|---|
| $T_{min}$ | DD | 3.0 ± 0.6 | 71 | 5.36 | $<10^{-6}$ |
| | FR | 4.8 ± 0.9 | 24 | 5.38 | $<10^{-4}$ |
| | IN | 3.4 ± 0.6 | 51 | 5.35 | $<10^{-5}$ |
| | SG | 0.3 ± 0.4 | 86 | 0.62 | 0.54 |
| | SP | 3.1 ± 0.6 | 34 | 5.21 | $<10^{-5}$ |
| $T_{opt}$ | DD | -2.5 ± 0.4 | 79 | -5.71 | $<10^{-6}$ |
| | FR | -2.3 ± 1.0 | 32 | -2.34 | 0.027 |
| | IN | -3.0 ± 0.6 | 70 | -5.36 | $<10^{-6}$ |
| | SG | -0.3 ± 0.3 | 114 | -0.93 | 0.36 |
| | SP | -2.1 ± 0.7 | 34 | -2.88 | 0.007 |
| $T_{max}$ | DD | -4.7 ± 0.7 | 58 | -6.62 | $<10^{-7}$ |
| | FR | -4.2 ± 1.2 | 25 | -3.43 | 0.002 |
| | IN | -3.7 ± 0.7 | 54 | -5.50 | $<10^{-5}$ |
| | SG | 0.0 ± 0.3 | 90 | 0.10 | 0.92 |
| | SP | -3.5 ± 0.8 | 31 | -4.3 | $<10^{-3}$ |
| $T_{range}$ | DD | -7.6 ± 1.0 | 53 | -7.76 | $<10^{-9}$ |
| | FR | -8.8 ± 1.5 | 22 | -5.76 | $<10^{-5}$ |
| | IN | -7.4 ± 1.1 | 43 | -6.80 | $<10^{-7}$ |
| | SG | -0.3 ± 0.7 | 81 | -0.50 | 0.62 |
| | SP | -6.4 ± 1.1 | 31 | -5.88 | $<10^{-5}$ |
| Skew | DD | -0.05 ± 0.02 | 46 | -2.35 | 0.023 |
| | FR | -0.09 ± 0.05 | 15 | -1.73 | 0.10 |
| | IN | -0.08 ± 0.03 | 37 | -3.02 | 0.005 |
| | SG | 0.00 ± 0.01 | 76 | 0.17 | 0.87 |
| | SP | -0.06 ± 0.03 | 25 | -1.99 | 0.058 |

**Table S3.**

Correlation coefficients and RMSE (°C) among $T_{opt}$ values for different processes within species. Degrees of freedom are given in parentheses. P-values are not given due to multiplicity of tests. NA denotes insufficient data.

| Correlation coefficients | | | | | |
|---|---|---|---|---|---|
| $T_{opt}$ | DD | FR | GC | IN | SG |
| DD | - | - | - | - | - |
| FR | 0.33 (11) | - | - | - | - |
| GC | 0.64 (78) | 0.32 (31) | - | - | - |
| IN | 0.95 (57) | 0.32 (11) | 0.64 (69) | - | - |
| SG | 0.75 (48) | 0.20 (17) | 0.69 (113) | 0.72 (52) | - |
| SP | 0.80 (16) | NA | 0.66 (33) | 0.83 (19) | 0.81 (26) |

| RMSE | | | | | |
|---|---|---|---|---|---|
| $T_{opt}$ | DD | FR | GC | IN | SG |
| DD | - | - | - | - | - |
| FR | 5.78 | - | - | - | - |
| GC | 4.70 | 6.09 | - | - | - |
| IN | 1.56 | 5.36 | 5.51 | - | - |
| SG | 4.08 | 6.69 | 3.20 | 3.95 | - |
| SP | 3.79 | NA | 4.66 | 3.36 | 4.75 |

**Table S4.**

Correlation coefficients among $T_{range}$ values for different processes within species. Degrees of freedom are given in parentheses. P-values are not given due to multiplicity of tests. NA denotes insufficient data.

| $T_{range}$ | DD | FR | GC | IN | SG |
|---|---|---|---|---|---|
| DD | - | - | - | - | - |
| FR | 0.12 (8) | - | - | - | - |
| GC | 0.11 (52) | 0.12 (21) | - | - | - |
| IN | 0.94 (42) | 0.21 (7) | 0.11 (42) | - | - |
| SG | 0.40 (33) | 0.27 (11) | 0.41 (80) | 0.27 (38) | - |
| SP | 0.46 (13) | NA | 0.42 (30) | 0.21 (16) | 0.43 (21) |

**Table S5.**

Correlation between abiotic niche breadth and biotic niche breadth. Abiotic niche breadth was estimated as $T_{range}$ or $T_{range(50\%)}$, and biotic niche breadth as $\log_{10+1}$-transformed host phylogenetic diversity calculated from either the processed or unprocessed host phylogeny. Parameter estimate, 95% confidence intervals, and t-test statistics vs. zero correlation are given.

| **$T_{range}$ / processed host phylogeny** | | | | | |
|---|---|---|---|---|---|
| **Process** | **Cor** | **95% CI** | **t** | **df** | **p** |
| DD | 0.200 | -0.077, 0.449 | 1.44 | 50 | 0.15 |
| FR | 0.000 | -0.481, 0.480 | 0.00 | 15 | 1.00 |
| GC | 0.128 | -0.018, 0.268 | 1.73 | 181 | 0.09 |
| IN | 0.008 | -0.265, 0.280 | 0.06 | 50 | 0.95 |
| SG | 0.198 | 0.002, 0.380 | 2.00 | 98 | 0.05 |
| SP | -0.108 | -0.490, 0.309 | -0.51 | 19 | 0.61 |

| **$T_{range}$ / unprocessed host phylogeny** | | | | | |
|---|---|---|---|---|---|
| **Process** | **Cor** | **95% CI** | **t** | **df** | **p** |
| DD | 0.218 | -0.058, 0.463 | 1.58 | 50 | 0.12 |
| FR | -0.100 | -0.554, 0.400 | -0.39 | 15 | 0.70 |
| GC | 0.140 | -0.001, 0.281 | 1.88 | 177 | 0.06 |
| IN | 0.090 | -0.188, 0.354 | 0.64 | 50 | 0.53 |
| SG | 0.195 | 0.003, 0.377 | 1.95 | 97 | 0.05 |
| SP | -0.177 | -0.542, 0.244 | -0.84 | 22 | 0.41 |

| **$T_{range(50\%)}$ / processed host phylogeny** | | | | | |
|---|---|---|---|---|---|
| **Process** | **Cor** | **95% CI** | **t** | **df** | **p** |
| DD | 0.071 | -0.230, 0.361 | 0.46 | 42 | 0.64 |
| FR | -0.081 | -0.626, 0.517 | -0.26 | 10 | 0.80 |
| GC | -0.007 | -0.154, 0.141 | -0.08 | 174 | 0.93 |
| IN | -0.108 | -0.400, 0.203 | -0.69 | 40 | 0.50 |
| SG | 0.015 | -0.187, 0.216 | 0.15 | 93 | 0.88 |
| SP | -0.186 | -0.572, 0.267 | -0.82 | 19 | 0.42 |

| **$T_{range(50\%)}$ / unprocessed host phylogeny** | | | | | |
|---|---|---|---|---|---|
| **Process** | **Cor** | **95% CI** | **t** | **df** | **p** |
| DD | 0.089 | -0.214, 0.376 | 0.58 | 42 | 0.57 |
| FR | -0.041 | -0.601, 0.546 | -0.13 | 10 | 0.90 |
| GC | -0.015 | -0.164, 0.135 | -0.20 | 170 | 0.84 |
| IN | -0.095 | -0.387, 0.216 | -0.60 | 40 | 0.55 |
| SG | -0.012 | -0.214, 0.191 | -0.11 | 92 | 0.91 |
| SP | 0.000 | -0.432, 0.431 | 0.00 | 19 | 1.00 |

**Table S6.**

Species names updated in the PlantWise database to ensure correct matching to the Togashi database. Species names were updated according to either the SFD or Mycobank. Species names were only updated in the PlantWise database if they were present in the Togashi dataset.

| Species name in the PlantWise database | Updated species name |
|---|---|
| *Acremonium strictum* | *Sarocladium strictum* |
| *Ascochyta gossypii* | *Ascochyta gossypiicola* |
| *Ascochyta pisi* | *Didymella pisi* |
| *Botryosphaeria obtusa* | *Peyronellaea obtusa* |
| *Botryosphaeria ribis* | *Neofusicoccum ribis* |
| *Cochliobolus heterostrophus* | *Bipolaris maydis* |
| *Cochliobolus miyabeanus* | *Bipolaris oryzae* |
| *Cochliobolus sativus* | *Bipolaris sorokiniana* |
| *Diaporthe phaseolorum* | *Phomopsis phaseoli* |
| *Diaporthe vaccinii* | *Phomopsis vaccinii* |
| *Fusarium coeruleum* | *Fusarium caeruleum* |
| *Fusicladium effusum* | *Venturia effusa* |
| *Gibberella avenacea* | *Fusarium avenaceum* |
| *Gibberella fujikuroi* | *Fusarium fujikuroi* |
| *Gibberella zeae* | *Fusarium graminearum* |
| *Glomerella cingulata* | *Colletotrichum gloeosporioides* |
| *Glomerella tucumanensis* | *Colletotrichum falcatum* |
| *Guignardia bidwellii* | *Phyllosticta ampelicida* |
| *Guignardia citricarpa* | *Phyllosticta citricarpa* |
| *Haematonectria haematococca* | *Fusarium solani* |
| *Hyaloperonospora parasitica* | *peronospora parasitica* |
| *Khuskia oryzae* | *Nigrospora oryzae* |
| *Leptosphaeria coniothyrium* | *Paraconiothyrium fuckelii* |
| *Magnaporthe salvinii* | *Nakataea oryzae* |
| *Mycosphaerella graminicola* | *Zymoseptoria tritici* |
| *Mycosphaerella pinodes* | *Didymella pinodes* |
| *Nectria coccinea* | *Neonectria coccinea* |
| *Olpidium brassicae* | *Olpidiaster brassicae* |
| *Ophiostoma piceae* | *Pesotum piceae* |
| *Passalora fulva* | *Fulvia fulva* |
| *Peronospora hyoscyami* f.sp. *Tabacina* | *Peronospora hyoscyami* |
| *Phellinus robustus* | *Fomitiporia robusta* |
| *Phoma destructiva* | *Remotididymella destructiva* |
| *Phoma pinodella* | *Didymella pinodella* |
| *Phoma tracheiphila* | *Plenodomus tracheiphilus* |
| *Phytophthora erythroseptica* var. *erythroseptica* | *Phytophthora erythroseptica* |
| *Pleospora herbarum* | *Stemphylium vesicarium* |
| *Pythium debaryanum* | *Globisporangium debaryanum* |
| *Pythium splendens* | *Globisporangium splendens* |
| *Pythium vexans* | *Phytopythium vexans* |
| *Rhizoctonia zeae* | *Waitea circinata* |
| *Setosphaeria turcica* | *Exserohilum turcicum* |
| *Sphacelotheca reiliana* | *Sporisorium reilianum* |
| *Thanatephorus cucumeris* | *Rhizoctonia solani* |
| *Ustilago nuda* f.sp. *Hordei* | *Ustilago nuda* |

**Table S7.**

Corrections to host species names in the PlantWise database to ensure correct phylogeny construction. Where a species or genus could not be identified in S.Phylo.Maker (NA), these were removed from phylogeny construction.

| **Host species name (PlantWise database)** | **Corrected host species name (S.Phylo.Maker)** |
|---|---|
| *Abelmoschus esculentus* | NA |
| *Acroptilon repens* | *Rhaponticum repens* |
| *Chamomilla recutita* | *Matricaria chamomilla* |
| *Chamomilla recutita* | *Matricaria chamomilla* |
| *Cheiranthus* spp. | *Erysimum* spp. |
| *Coleus* spp. | NA |
| *Cuprocyparis leylandii* | NA |
| *Cyphomandra betacea* | NA |
| *Dizygotheca* spp. | *Schefflera* spp. |
| *Elettaria* spp. | NA |
| *Elettaria cardamomum* | NA |
| *Gloriosa rothschildiana* | NA |
| *Pascopyrum smithii* | *Elymus smithii* |
| *Pharbitis nil* | *Ipomoea nil* |
| *Pharbitis purpurea* | *Ipomoea purpurea* |
| *Poncirus* spp. | *Citrus* spp. |
| *Samanea saman* | *Albizia saman* |
| *Scindapsus* spp. | *Epipremnum* spp. |
| *Ullucus tuberosus* | NA |
| *Xanthocyparis nootkatensis* | *Cupressus nootkatensis* |

**References and Notes:**

38. Dryad. Deposited dataset.
39. Togashi, K. Biological Characters of Plant Pathogen Temperature relations. Meibundo. Tokyo.
40. Martin, Frank N., et al. "Identification and detection of Phytophthora: reviewing our progress, identifying our needs." *Plant Disease* 96.8 (2012): 1080-1103.
41. Magarey, R. D., T. B. Sutton, and C. L. Thayer. "A simple generic infection model for foliar fungal plant pathogens." *Phytopathology* 95.1 (2005): 92-100.
42. R Core Team (2019). R: A language and environment for statistical computing. R Foundation for Statistical Computing, Vienna, Austria. URL https://www.R-project.org/.
43. Yan, Weikai, and L. A. Hunt. "An equation for modelling the temperature response of plants using only the cardinal temperatures." *Annals of Botany* 84.5 (1999): 607-614.
44. Pinheiro J, Bates D, DebRoy S, Sarkar D, R Core Team (2019). *nlme: Linear and Nonlinear Mixed Effects Models*. R package version 3.1-141, https://CRAN.R-project.org/package=nlme.
45. Qian, Hong, and Yi Jin. "An updated megaphylogeny of plants, a tool for generating plant phylogenies and an analysis of phylogenetic community structure." *Journal of Plant Ecology* 9.2 (2016): 233-239.
46. S.W. Kembel, P.D. Cowan, M.R. Helmus, W.K. Cornwell, H. Morlon, D.D. Ackerly, S.P. Blomberg, and C.O. Webb. 2010. Picante: R tools for integrating phylogenies and ecology. Bioinformatics 26:1463-1464.
47. Faith, Daniel P. "Conservation evaluation and phylogenetic diversity." *Biological conservation* 61.1 (1992): 1-10.
48. Revell, Liam J. "phytools: an R package for phylogenetic comparative biology (and other things)." *Methods in Ecology and Evolution* 3.2 (2012): 217-223.
49. Pasiecznik, N. M., et al. "CABI/EPPO distribution maps of plant pests and plant diseases and their important role in plant quarantine." *Eppo Bulletin* 35.1 (2005): 1-7.
50. Bebber, Daniel P., Mark AT Ramotowski, and Sarah J. Gurr. "Crop pests and pathogens move polewards in a warming world." *Nature climate change* 3.11 (2013): 985.
51. Goslee, Sarah C., and Dean L. Urban. "The ecodist package for dissimilarity-based analysis of ecological data." *Journal of Statistical Software* 22.7 (2007): 1-19.
52. Balbuena, Juan Antonio, Raúl Míguez-Lozano, and Isabel Blasco-Costa. "PACo: a novel procrustes application to cophylogenetic analysis." *PloS one* 8.4 (2013): e61048.
53. Hutchinson, Matthew C., et al. "paco: implementing Procrustean Approach to Cophylogeny in R." *Methods in Ecology and Evolution* 8.8 (2017): 932-940.
